# An immunosuppression imaging gauge prospectively predicts metastatic progression and immunotherapy resistance in breast cancer

**DOI:** 10.64898/2026.09.22.753629

**Authors:** Yuwei Zhang, Jing Wang

**Affiliations:** Chemical and Biological Engineering Department, Iowa State University, Ames, IA, 50011

## Abstract

Systemic immunosuppression promotes metastatic progression and immunotherapy resistance, yet reliable biomarkers and noninvasive tools for longitudinal monitoring remain limited. Building on our finding that CXCL1⁺ neutrophil abundance in major organs inversely correlates with breast cancer–induced systemic immunosuppression, we developed an imaging-based immunosuppression gauge using CXCR2-expressing cells that mimic CXCL1⁺ neutrophil trafficking. Following intravenous administration, the biodistribution of engineered CXCR2-overexpressing mesenchymal stem cells or healthy donor neutrophils was imaged at 24 h, with greater organ accumulation indicating lower systemic immunosuppression. These living, distributed biosensors integrate chemokine signals across multiple organs and convert host immune status into an imageable biodistribution pattern. The gauge distinguished tumor aggressiveness at early disease stages, predicted metastatic risk before overt metastatic outgrowth, identified chemoimmunotherapy responders, and informed treatment selection through longitudinal immune monitoring. Mechanistically, nonaggressive tumors failed to induce substantial systemic immunosuppression and were effectively controlled by surgery alone, whereas aggressive tumors continued to drive immunosuppression after resection. Neoadjuvant chemoimmunotherapy produced superior outcomes to adjuvant treatment by intervening before profound immunosuppression developed and more effectively restoring antitumor immunity. Across cancer models, long-term survivors consistently showed resolution of systemic immunosuppression and restoration of immune activity to healthy-host levels. Thus, imaging systemic immunosuppression provides an early, longitudinal indicator of metastatic risk, therapeutic response, and treatment efficacy.

**Statement of significance:** A living-cell imaging gauge converts systemic immune status into an imageable signal, enabling early prediction of metastatic progression and immunotherapy response while revealing systemic immunosuppression as an actionable determinant of treatment outcome.

## Introduction

Systemic immunosuppression, characterized by expansion of suppressive myeloid, impaired T-cell function, and reduced elimination of tumor cells, is a hallmark of advanced cancer. It precedes and promotes metastatic outgrowth and contributes to immunotherapy resistance (1–3). Quantifying the magnitude of immunosuppression and monitoring its dynamic changes during cancer progression or therapy could therefore inform metastatic risk and therapeutic resistance earlier than conventional imaging approaches such as computed tomography (CT), which typically detect established tumor masses only after they have reached millimeter-scale size and contain millions of tumor cells (4). Currently, one of the most direct methods to quantify immunosuppressive activity in preclinical models is to isolate myeloid cells and T cells from tumor-bearing animals and perform ex vivo T-cell proliferation assays. However, this approach is terminal, labor-intensive, and unable to continuously monitor immune changes within the same host over time (5). Clinically, liquid biopsy–based approaches have been explored to estimate cancer-associated immunosuppression by measuring blood-cell ratios such as the neutrophil-to-lymphocyte ratio; quantifying circulating suppressive immune populations such as myeloid-derived suppressor cells (MDSCs); assessing immune cell suppressive function; and profiling circulating cytokines, selected immune biomarkers, or transcriptomic signatures (6–12). However, these approaches could be either technically complex or insufficiently accurate and timely for predicting metastatic risk and therapeutic resistance. Additionally, some readouts such as cytokine levels can be influenced by infection, inflammation, and other non-cancer-related pathological conditions. These limitations highlight the need for a noninvasive, specific, and longitudinally applicable method to monitor cancer-associated systemic immunosuppression. Such a method would enable clinical assessment of metastatic risk and therapy resistance, while also supporting preclinical studies of metastatic mechanisms and therapeutic evaluation.

The major challenge in developing such an immunosuppression gauge is the identification of disease-specific biomarkers. We recently reported that CXCL1⁺ neutrophils, a subset characterized by elevated CXCL1 production and migration toward CXCL1 and other CXCR2 ligands, play a key role in promoting pro-inflammatory and antitumor responses in breast cancer. A high abundance of these cells in major organs indicates a low level of systemic immunosuppression and is therefore associated with reduced metastatic risk and favorable prognosis (13). Consistently, clinical data suggest that chemoattractants for these cells serve as favorable prognostic markers in breast cancer (14). Here, we harnessed CXCL1⁺ neutrophils as an inverse biomarker of cancer-induced systemic immunosuppression and developed cell mimics that overexpress CXCR2 and therefore exhibit similar in vivo trafficking patterns. By tracking the biodistribution of intravenously injected CXCL1⁺ neutrophil mimics and comparing it with their distribution in healthy hosts, we can quantify systemic immunosuppression induced by breast cancer. These injected cells function as living distributed biosensors by integrating spatial chemokine signals across multiple organs and converting the host immune state into an imageable biodistribution pattern. Two types of cellular probes were evaluated. The first consists of mesenchymal stem cells (MSCs) engineered to stably overexpress CXCR2. MSCs have been widely used to deliver growth factors and cytokines, exhibiting a favorable safety profile in preclinical models (15). They are easy and cost-effective to prepare, making them suitable for animal studies. The second consists of healthy donor-derived neutrophils that endogenously express high levels of CXCR2. Clinically, neutrophil transfusion has been used to treat neutropenia (16), and neutrophil-based imaging has been applied to identify sites of inflammation or infection, supporting their translational potential (17,18). To validate this imaging-based immunosuppression gauge for cancer diagnosis and therapy monitoring, we evaluated its ability to distinguish tumor aggressiveness and predict prognosis at early disease stages, compared its timeliness in detecting metastatic risk with conventional tumor-imaging technologies, and assessed its accuracy in providing early indications of therapy resistance during treatment response monitoring.

## Results

### CXCR2⁺ cell accumulation in major organs quantifies systemic immunosuppression comparably to ex vivo T-cell suppression assays and enables early discrimination of breast tumor aggressiveness

We first used breast cancer preclinical models to validate the immunosuppression gauge, which uses the frequency of CXCL1⁺ neutrophils in major organs as an inverse biomarker of cancer-induced systemic immunosuppression. The highly aggressive murine breast cancer 4T1 and its two well-characterized sublines—moderately aggressive 4T07 and non-aggressive 67NR— differ in their ability to establish systemic immunosuppressive host environments (19,20). Consistent with their metastatic potential, increasing tumor aggressiveness was associated with higher frequencies of immunosuppressive populations, including myeloid-derived suppressor cells (MDSCs) and regulatory T cells (Tregs), and lower frequencies of immune-active populations, including Th1 cells, Th17 cells, activated CD8⁺ T cells, and activated natural killer (NK) cells (**Fig. 1A**). Flow cytometric gating strategies for these immune populations are shown in **Fig. S1**. These phenotypic differences were accompanied by corresponding differences in functional immunosuppression. At a primary tumor size of ∼1000 mm³, lung-derived Gr1⁺ myeloid cells from 4T07- and 4T1-bearing BALB/c mice significantly suppressed T-cell proliferation, whereas those isolated from 67NR-bearing mice had no significant suppressive effect (**Fig. 1B**). Because immunosuppression in distant organs can create a permissive environment for metastatic colonization and outgrowth (21–23), these functional differences have direct relevance to metastatic progression. Following intracardiac injection, 4T1 cells seeded the lungs of both 4T1- and 67NR-bearing mice but developed into secondary tumors only in the immunosuppressive lungs of 4T1-bearing mice, whereas they were eliminated from the immune-active lungs of 67NR-bearing mice (13). We therefore used these well-characterized models to determine whether our immunosuppression gauge could quantitatively measure tumor-induced systemic immunosuppression in vivo.

**Fig. 1.**
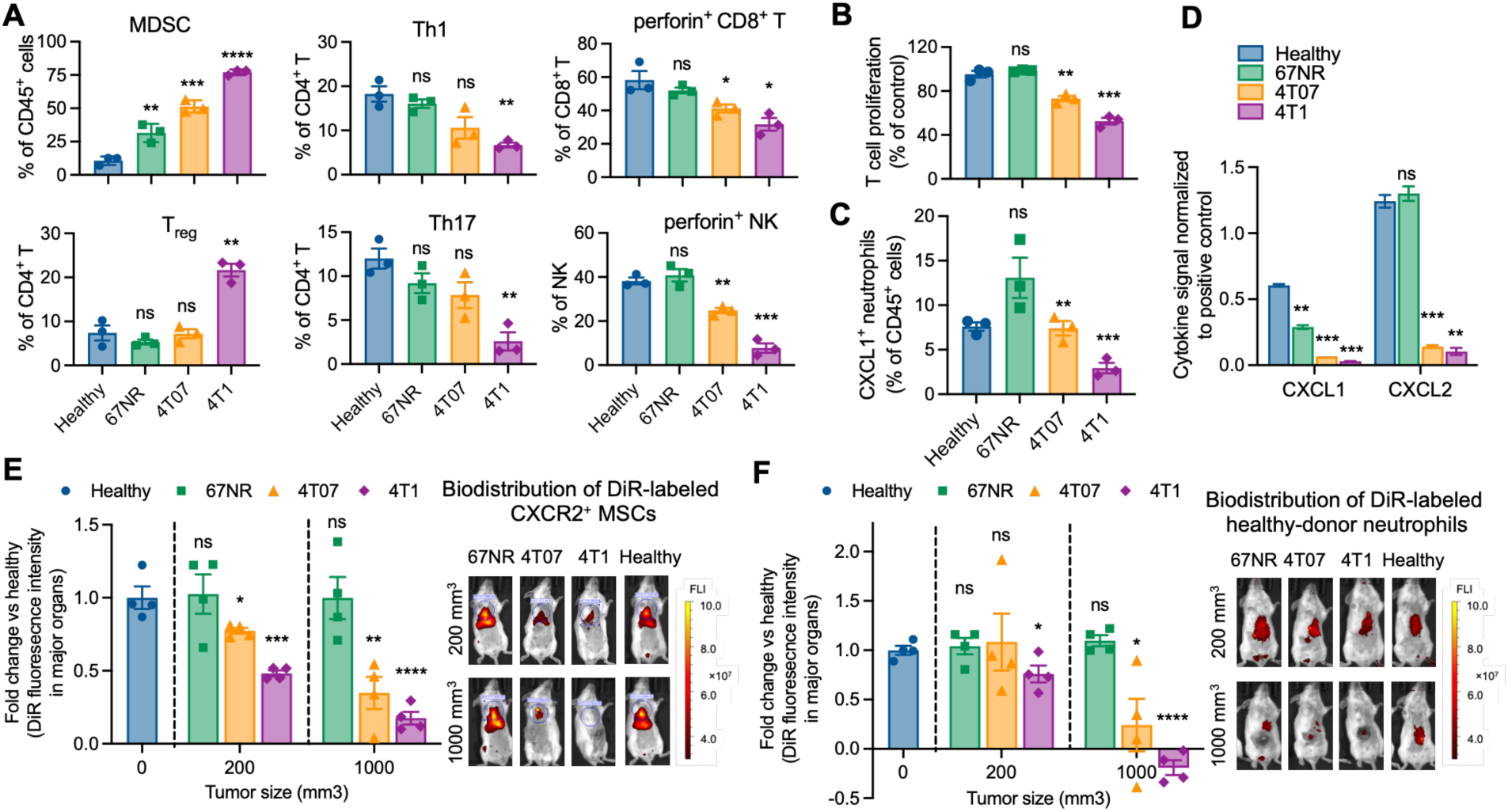
The immunosuppression gauge monitors CXCR2⁺ cell trafficking and quantifies systemic immunosuppression and enables early discrimination of tumor aggressiveness. (A) Frequencies of immunosuppressive and immune-active cell populations in the lungs of tumor-free healthy BALB/c mice or mice bearing 67NR, 4T07, or 4T1 tumors. (B) Suppressive activity of lung-derived Gr1⁺ myeloid cells on T-cell proliferation. (C, D) Frequencies of CXCL1⁺ neutrophils and the abundance of their chemoattractants, including CXCL1 and CXCL2, in mouse lungs. For (A-D), tissues were from tumor-free mice or mice bearing ∼1000 mm³ primary tumors. Tumor aggressiveness followed the order 67NR < 4T07 < 4T1. (E, F) Imaging of DiR-labeled CXCR2⁺ MSCs or healthy donor neutrophils in tumor-free BALB/c mice and tumor-bearing mice. The distribution of DiR-labeled cells was assessed by IVIS imaging 24 hours after intravenous injection. Regions of interest (ROI) used to quantify DiR fluorescence of CXCR2^+^ MSCs and healthy donor neutrophils in BALB/c mice are shown in Fig. S3 and Fig S4, respectively. CXCR2⁺ cell accumulation mirrored CXCL1⁺ neutrophil abundance and was inversely associated with systemic immunosuppression and tumor aggressiveness. MDSC: myeloid-derived suppressive cell. Th: helper T cell. NK: natural killer. MSC: mesenchymal stem cell. ns: not significant. *p < 0.05, **p < 0.01, and ***p < 0.001, and ***p < 0.0001. P values in (A–F) were calculated relative to the healthy control group.

Across these breast cancer preclinical models, the degree of systemic immunosuppression was inversely associated with the abundance of CXCL1⁺ neutrophils, an immune-active neutrophil subset with potent antitumor activity, and with levels of their chemoattractants, including CXCL1 and CXCL2, in major organs (**Fig. 1C, 1D**). We therefore sought to use the abundance of CXCL1⁺ neutrophils in major organs as an inverse indicator of tumor-induced systemic immunosuppression. In our previous studies, we found that the in vivo trafficking of CXCL1⁺ neutrophils was regulated by the CXCR2–ligand axis (13). Based on this mechanism, we developed two CXCR2⁺ cell populations as surrogates for CXCL1⁺ neutrophils: mesenchymal stem cells (MSCs) engineered to stably overexpress CXCR2 by lentiviral transfection and neutrophils isolated from healthy donor mice by magnetic-activated cell sorting (MACS). Engineered CXCR2⁺ MSCs provide a reproducible and cost-effective platform for preclinical studies, whereas healthy donor neutrophils offer greater translational relevance for potential clinical applications. Flow cytometric analysis confirmed elevated surface CXCR2 expression on engineered CXCR2⁺ MSCs and healthy donor neutrophils relative to their respective controls, including nonengineered MSCs and the non-neutrophil fraction obtained during MACS (**Fig. S2**). To evaluate their trafficking in vivo, CXCR2⁺ MSCs or healthy donor neutrophils were labeled with DiR dyes, intravenously injected into tumor-bearing or healthy mice, and imaged 24 hours later. DiR fluorescence in major organs, including the lungs and liver, was quantified and normalized to the corresponding signals in healthy mice to provide a quantitative measure of host immune status. The regions of interest (ROI) used to define major organs and the reference ranges of DiR fluorescence observed in healthy BALB/c mice are shown in **Fig. S3** and **Fig. S4** for CXCR2⁺ MSCs and healthy donor neutrophils, respectively.

Both CXCR2⁺ MSCs and healthy donor neutrophils reproduced the organ-level trafficking pattern of endogenous CXCL1⁺ neutrophils (**Fig. 1E, 1F**, **1C**). Their accumulation in major organs correlated with the abundance of CXCR2 ligands, including CXCL1 and CXCL2, and progressively decreased with increasing systemic immunosuppression (**Fig. 1D**). In contrast, CXCR2-negative cells, including nonengineered MSCs, failed to preferentially accumulate in organs enriched in CXCR2 ligands after intravenous injection (**Fig. S5**). These findings indicate that the trafficking of CXCR2⁺ MSCs and healthy donor neutrophils, similar to that of endogenous CXCL1⁺ neutrophils, is governed by the CXCR2–ligand axis. Thus, the capacity of major organs to recruit CXCR2⁺ cells provides a quantitative in vivo readout of systemic immune status. CXCR2⁺ MSCs exhibited relatively uniform CXCR2 expression and consistent biodistribution among healthy mice. In contrast, healthy donor neutrophils showed greater donor-to-donor variability in trafficking, likely reflecting differences in CXCR2 expression among donors (**Figs. S3, S4**). Therefore, for donor neutrophil-based imaging, tumor-free healthy mice were imaged alongside tumor-bearing mice using neutrophils from the same donor to provide a donor-matched reference. This additional control was not required for CXCR2⁺ MSC-based imaging because of their more uniform CXCR2 expression and reproducible biodistribution. Importantly, intravenously administered CXCR2⁺ MSCs and healthy donor neutrophils were cleared within 2– 5 days (**Fig. S6**), enabling repeated administration and longitudinal monitoring of host immune status at weekly intervals. Clearance was more rapid in highly immunosuppressive 4T1-bearing mice than in healthy controls.

We compared the immunosuppression gauge with the conventional ex vivo T-cell suppression assay. At a primary tumor size of ∼1000 mm³, T-cell suppression by lung-derived Gr1⁺ myeloid cells (**Fig. 1B**), CXCR2⁺ MSC imaging (**Fig. 1E**), and healthy donor neutrophil imaging (**Fig. 1F**) consistently showed that the capacity of the three tumor models to induce systemic immunosuppression paralleled their aggressiveness, following the order 4T1 > 4T07 > 67NR. These results establish CXCR2⁺ cell imaging as a noninvasive alternative for quantifying systemic immunosuppression without requiring tissue collection and ex vivo functional assays. Notably, CXCR2⁺ MSC imaging discriminated among these tumor models when primary tumors were only ∼200 mm³, an early disease stage at which significant T-cell suppressive activity had not yet developed (**Fig. 2E**). This earlier discrimination likely reflects the fact that changes in CXCL1⁺ neutrophil abundance precede the establishment of overt functional T-cell suppression. By using CXCL1⁺ neutrophil recruitment as an early indicator of tumor–host immune interactions, the CXCR2⁺ cell-based immunosuppression gauge can therefore detect differences in tumor-induced systemic immune status before conventional functional immunosuppression becomes apparent. This capability enables early discrimination of tumor aggressiveness and metastatic potential and may provide information relevant to therapeutic selection and prognosis.

**Fig. 2.**
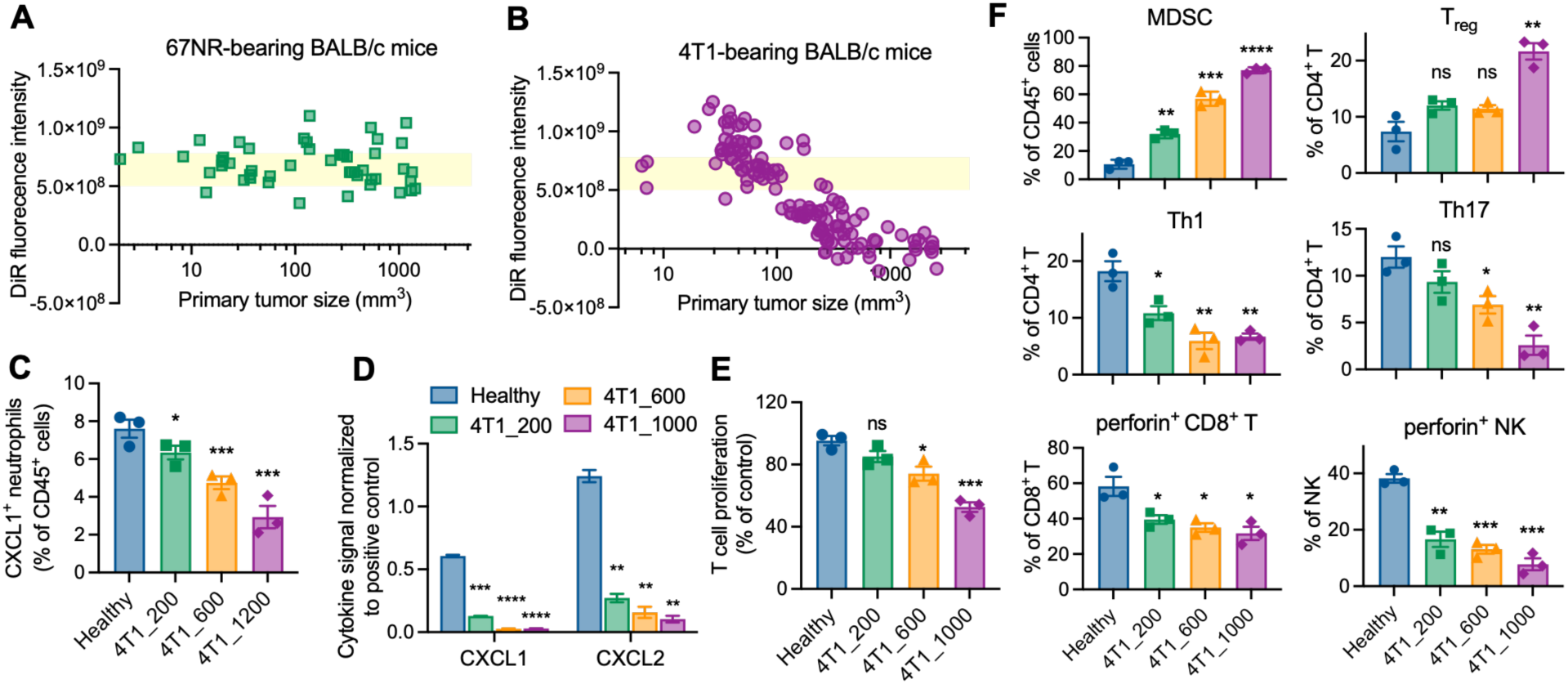
The immunosuppression gauge captures tumor aggressiveness-dependent immune dynamics and predicts metastatic risk before overt metastatic outgrowth. BALB/c mice bearing 67NR or 4T1 tumors were longitudinally monitored by weekly intravenous injection and imaging of DiR-labeled CXCR2⁺ MSCs. (A) In mice bearing non-aggressive 67NR tumors, lung accumulation of CXCR2⁺ MSCs remained similar to that in tumor-free mice across cancer progression, indicating preserved systemic immune activity and minimal immunosuppression. (B) In mice bearing highly aggressive 4T1 tumors, the immunosuppression gauge classified host immune progression into three stages: early immune activation, developing immunosuppression, and established functional immunosuppression. Yellow bars in (A, B) indicate the DiR fluorescence intensity range measured in healthy BALB/c, as identified in Fig S3. (C, D) Abundance of CXCL1⁺ neutrophils and their chemoattractants in the lungs of 4T1-bearing mice decreased as primary tumors grew. (E) Functional T-cell suppression by lung-derived Gr1^+^ myeloid cells became detectable when 4T1 tumors were > 600 mm^3^. (F) Frequencies of immunosuppressive and immune-active cell populations in the lungs of 4T1-bearing mice with varying tumor sizes. Numbers in (C-F) indicate primary tumor size in mm³. ns: not significant. *p < 0.05, **p < 0.01, ***p < 0.001, and ***p < 0.0001. P values in (C–F) were calculated relative to the healthy control group.

Collectively, these results establish CXCR2⁺ MSCs and healthy donor neutrophils as functional surrogates for endogenous CXCL1⁺ neutrophil trafficking and demonstrate that their recruitment to major organs can be used to quantitatively and longitudinally monitor systemic immune status in breast cancer models. Importantly, by detecting early changes in CXCL1⁺ neutrophil recruitment, the immunosuppression gauge distinguishes differences in tumor aggressiveness and metastatic potential before the development of overt functional immunosuppression.

### The immunosuppression gauge captures distinct immune dynamics in non-aggressive versus aggressive breast cancers and predicts metastatic risk before overt metastatic outgrowth

We next applied CXCR2⁺ MSC imaging as a longitudinal gauge of systemic immunosuppression during cancer progression. BALB/c mice were orthotopically inoculated with 67NR or 4T1 tumor cells and subsequently received weekly intravenous administration of DiR-labeled CXCR2⁺ MSCs followed by imaging until the humane endpoint. Endpoints were reached because of excessive primary tumor growth in 67NR-bearing mice or metastasis-associated distress in 4T1-bearing mice (**Fig. 3A**). Longitudinal imaging revealed markedly different tumor– host immune trajectories between the non-aggressive 67NR model and the highly aggressive 4T1 model.

**Fig. 3.**
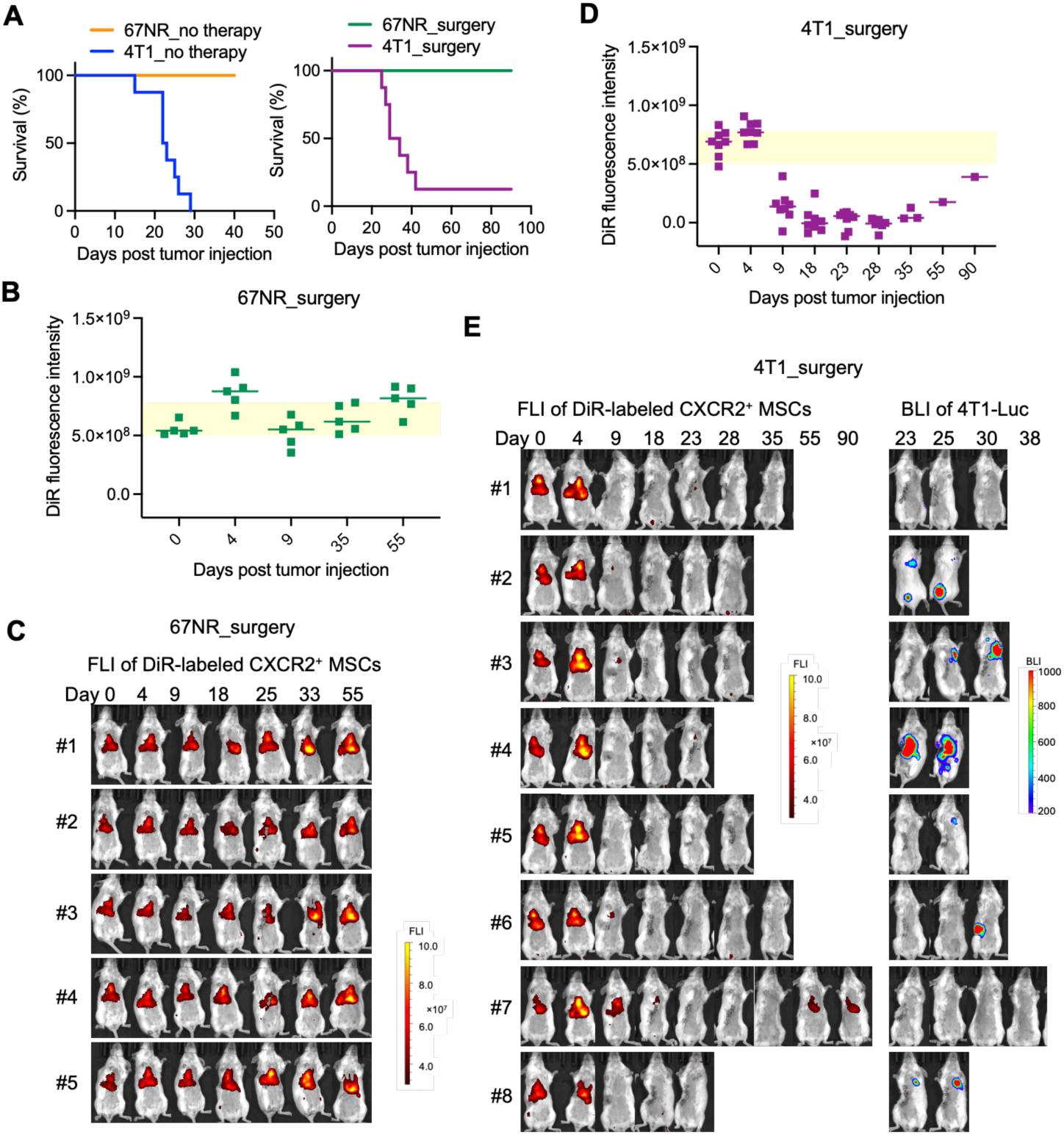
The immunosuppression gauge predicts tumor aggressiveness-dependent outcomes after surgery. (A) Kaplan–Meier survival curves of 67NR- and 4T1-bearing BALB/c mice without any treatment (left) or following surgical resection of the primary tumor (right). Untreated 67NR-bearing mice reached the experimental endpoint because of large primary tumor size. (B, C) Longitudinal CXCR2⁺ MSC imaging in 67NR-bearing mice after surgery, showing preserved major organ recruitment of CXCR2⁺ MSCs comparable to healthy mice. (D, E) Longitudinal CXCR2⁺ MSC imaging and bioluminescence-based metastasis monitoring in 4T1-bearing mice after surgery. Mice that eventually succumbed to metastases showed progressive loss of major organ CXCR2⁺ MSC recruitment followed by metastatic outgrowth. In contrast, the mouse that achieved long-term survival restored CXCR2⁺ MSC recruitment to major organs, indicating alleviation of systemic immunosuppression. Notably, mouse #1 died with metastases that did not show luciferase signals. Mouse #7 achieved long-term survival. The primary tumors were resected at ∼400 mm^3^, which was day 24 after tumor inoculation in 67NR-bearing mice and day 12 after tumor inoculation in 4T1-bearing mice. FLI: fluorescence imaging; BLI: bioluminescence imaging.

Mice bearing 67NR tumors maintained an immune-active systemic state comparable to that of healthy mice throughout cancer progression, irrespective of primary tumor size (**Fig. 2A**). Even when 67NR tumors reached ∼1,000 mm³, major organs remained enriched in CXCL1⁺ neutrophils and their chemoattractants (**Figs. 1C, 1D**) and retained higher frequencies of immune-active populations, including Th1 cells, Th17 cells, activated CD8⁺ T cells, and activated NK cells (**Fig. 1A**). Consistently, Gr-1⁺ myeloid cells isolated from 67NR-bearing mice at this late stage did not suppress T-cell proliferation (**Fig. 1B**). These findings indicate that 67NR tumors fail to establish substantial systemic immunosuppression despite continued primary tumor growth. Because an immunosuppressive distant organ environment facilitates metastatic outgrowth, preservation of this immune-active state may explain why intravenously introduced 4T1 cells that seeded the lungs of 67NR-bearing mice failed to develop into secondary tumors (13).

In contrast, 4T1 tumors induced a progressive transition from systemic immune activation to established immunosuppression. Based on longitudinal CXCR2⁺ MSC recruitment and complementary immune measurements, this progression could be divided into three stages: early immune activation, developing immunosuppression, and established functional immunosuppression (**Fig. 2B**). During the first stage, when 4T1 primary tumors were < ∼30 mm³, CXCR2⁺ MSC accumulation in major organs exceeded that observed in healthy mice, indicating transient systemic immune activation shortly after tumor establishment. This response may reflect an early host respond to newly implanted tumor cells, consistent with reports that transplanted tumors rapidly recruit innate immune cells and that early 4T1 tumors exhibit immune-active transcriptional features before progressive immunosuppression develops (24–27).

During the second stage, as 4T1 primary tumors increased from ∼30 to ∼580 mm³, the systemic immune environment progressively shifted toward immunosuppression (**Fig. 2B**). CXCR2⁺ MSC recruitment to major organs gradually declined with tumor growth, paralleling reductions in CXCL1⁺ neutrophils and their chemoattractants (**Figs. 2C, 2D**). Concurrently, immunosuppressive populations, including MDSCs and Tregs, increased, whereas Th1 cells, Th17 cells, activated CD8⁺ T cells, and activated NK cells declined (**Fig. 2F**). Despite these phenotypic changes, lung-derived Gr-1⁺ myeloid cells had not yet acquired significant T-cell-suppressive activity (**Fig. 2E**), indicating that this period represents a transitional phase in which immunosuppression is developing but is not yet functionally established. Our previous studies suggest that the premetastatic niche develops during this stage and becomes increasingly permissive to circulating 4T1 cells disseminating from the primary tumor (13). However, overt macrometastases remain undetectable by H&E staining or CT imaging at this stage (28). Thus, systemic immune conditioning and metastatic dissemination appear to precede detectable metastatic outgrowth.

During the third stage, when 4T1 primary tumors exceeded ∼580 mm³, major organs largely lost their ability to recruit intravenously administered CXCR2⁺ MSCs (**Fig. 2B**), consistent with depletion of chemotactic signals for CXCL1⁺ neutrophils (**Fig. 2D**). At this point, functional immunosuppression was established, as lung-derived Gr-1⁺ myeloid cells markedly suppressed ex vivo T-cell proliferation (**Fig. 2E**). This immune state coincided with rapid metastatic expansion. In our previous studies, the number of 4T1/tdTomato/Luc cells in the lungs increased from hundreds to millions as primary tumors grew from ∼600 to ∼1500 mm³, while lung metastases became detectable by H&E staining and CT imaging only when primary tumors reached ∼1200 mm³ (13,28).

Collectively, longitudinal CXCR2⁺ MSC imaging in 67NR- and 4T1-bearing BALB/c mice revealed a tumor aggressiveness-dependent progression in systemic immune status. Less aggressive 67NR tumors maintained an immune-active host environment, whereas highly aggressive 4T1 tumors drove a staged transition from early immune activation to developing immunosuppression and ultimately to established functional immunosuppression. Importantly, the immunosuppression gauge identified the onset of systemic immunosuppression permissive for metastatic outgrowth at a primary tumor volume of ∼580 mm³, well before lung macrometastases became detectable by conventional CT imaging at ∼1200 mm³ (28).

### The immunosuppression gauge predicts distinct outcome of surgery monotherapy for non-aggressive versus aggressive breast cancers

Distinct tumor–host interactions suggest that cancers with different aggressiveness require customized therapy strategies. We therefore used the immunosuppression gauge to monitor immune dynamics in 67NR- and 4T1-bearing mice after early-stage tumor resection surgery. This analysis revealed that surgical outcome was strongly associated with tumor aggressiveness and the extent of tumor-induced systemic immunosuppression. Mice were orthotopically inoculated with tumor cells on day 0 and underwent surgical resection when primary tumors reached ∼400 mm³. In 67NR-bearing mice, surgery alone achieved 100% long-term survival, and the hosts maintained an immune-active status comparable to healthy mice after surgery (**Figs. 3A, 3B, 3C**). Major organs remained enriched with CXCL1⁺ neutrophils and their chemoattractants, as well as immune cells with pro-inflammatory and antitumor phenotypes (**Fig. 4**, 67NR_surgery vs. healthy). These results indicate that, for hosts bearing non-aggressive tumors that do not induce systemic immunosuppression, early tumor resection is sufficient to prevent disease progression.

**Fig. 4.**
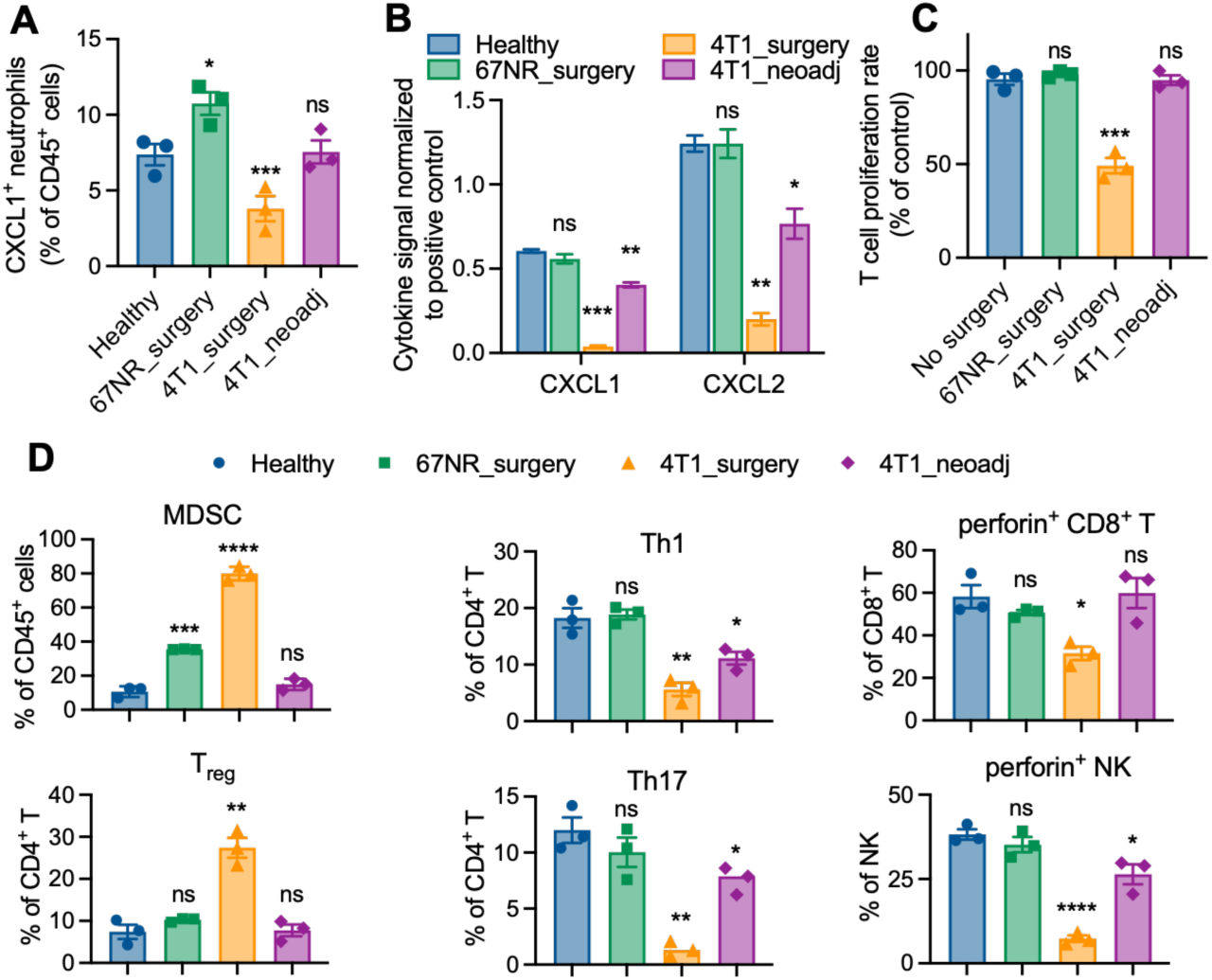
Therapy-responsive 67NR- or 4T1-bearing BALB/c mice alleviate immunosuppression and enhance CXCL1^+^ neutrophils in major organs. (A) Frequencies of CXCL1⁺ neutrophils, and (B) the abundance of their chemoattractants, including CXCL1 and CXCL2, and (C) T-cell suppressive ability of lung-derived Gr1⁺ myeloid cells, and (D) frequencies of immunosuppressive and immune-active cell populations in the lungs of healthy BALB/c mice or mice treated with surgery alone or neoadjuvant therapy. Primary tumors were resected on day 12 after tumor inoculation, when untreated mice had an average tumor volume of ∼400 mm³. Tissues were collected on day 90 after tumor inoculation from 67NR-bearing mice and from 4T1-bearing mice that received neoadjuvant therapy and achieved long-term survival. For 4T1-bearing mice treated with surgery alone, tissues were collected at the humane endpoint triggered by metastasis-associated distress. ns: not significant. *p < 0.05, **p < 0.01, ***p < 0.001, and ****p < 0.0001. P values were calculated relative to the healthy control group.

In contrast, 4T1-bearing mice continued to develop progressive immunosuppression after early-stage surgery, and 90% of mice eventually succumbed to distant metastases (**Figs. 3A, 3D, 3E**). Because 4T1 tumor cells expressed a luciferase reporter in this study, metastatic progression could be longitudinally monitored by bioluminescence imaging after surgery. In mice that failed to benefit from surgery, IVIS-detectable metastases emerged within 7-10 days after major organs completely lost the ability to recruit intravenously injected CXCR2⁺ MSCs, marking the onset of functional immunosuppression and metastatic outgrowth. Consistently, immune profiling showed that the lungs of post-surgical 4T1-bearing mice remained immunosuppressive and pro-tumor immediately before reaching the humane endpoint due to metastasis-associated distress (**Fig. 4**, 4T1_surgery vs. healthy). Notably, the only 4T1-bearing mouse that achieved metastasis-free long-term survival after surgery recovered the ability of recruiting CXCR2⁺ MSCs to major organs, indicating alleviation of systemic immunosuppression and restoration of antitumor immunity (**Figs. 3D, 3E**, mouse #7).

Together, these findings suggest that surgery may be sufficient to achieve durable disease control when breast tumors are less aggressive and systemic antitumor immunity remains intact. In contrast, surgery alone may be inadequate for aggressive, high-risk tumors even when the primary tumor is resected at an early stage, because removal of the primary tumor does not necessarily reverse systemic immunosuppression or prevent subsequent metastatic outgrowth. Consistent with these findings, clinical trials including PRIME II trial and CALGB 9343, have shown that patients with biologically favorable, low-risk breast cancers have a low risk of distant disease progression following complete resection, although adjuvant therapy can further reduce local recurrence (29,30). Conversely, trials including NSABP B-13, CREATE-X, KEYNOTE-522, and HERA have demonstrated substantial residual recurrence risk in biologically aggressive or high-risk early-stage breast cancer despite surgical resection and have shown that effective systemic adjuvant or perioperative therapy can improve disease control and, in selected settings, overall survival (31–34). These findings underscore the importance of systemic therapy for controlling occult disseminated disease in high-risk aggressive breast cancer.

### The immunosuppression gauge distinguishes immunotherapy resistance before detectable metastatic outgrowth in aggressive breast cancer

We next tested whether perioperative chemoimmunotherapy could reverse the immunosuppressive trajectory in 4T1-bearing mice. Neoadjuvant and adjuvant chemoimmunotherapy, consisting of two doses of cisplatin and four doses of anti-PD-1 administered before or after early-stage tumor resection, achieved long-term survival rates of 70% and 20%, respectively (**Fig. 5A**). Notably, neoadjuvant therapy significantly reduced primary tumor size at the time of surgery compared with surgery alone or adjuvant therapy (**Fig. S7**). The superior efficacy of neoadjuvant over adjuvant chemoimmunotherapy in eradicating metastasis diseases in the 4T1 model is consistent with previous reports (35,36). Using longitudinal CXCR2⁺ MSC imaging, we further examined how treatment timing influenced systemic immune recovery and whether the immunosuppression gauge could distinguish responders from nonresponders and predict long-term outcome.

**Fig. 5.**
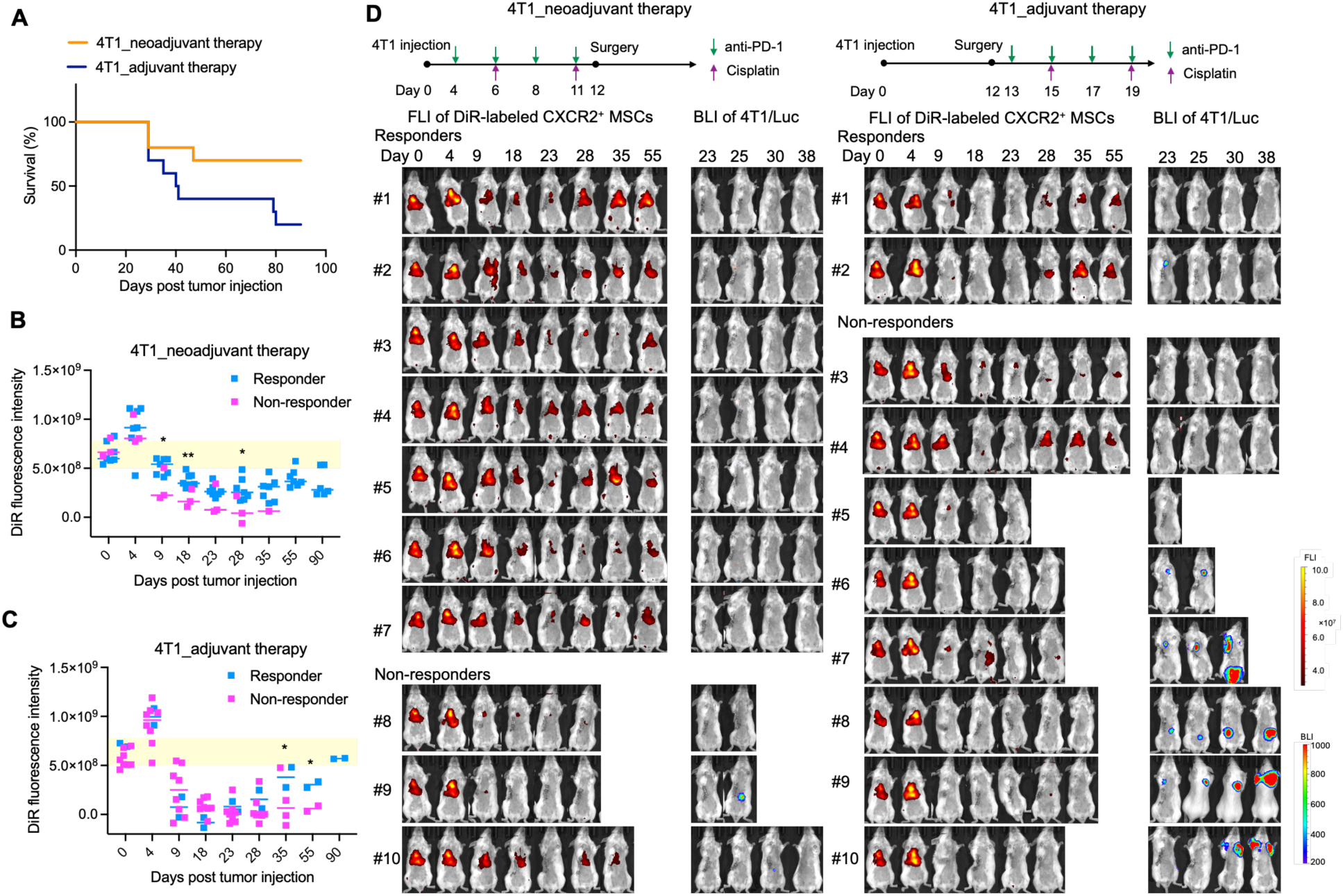
The immunosuppression gauge predicts neoadjuvant and adjuvant chemoimmunotherapy response before metastatic outgrowth. 4T1-bearing mice underwent early-stage surgery with perioperative cisplatin and anti-PD-1 administered as neoadjuvant or adjuvant therapy. (A) Kaplan–Meier survival curves comparing treatment outcomes. (B, C) Longitudinal CXCR2⁺ MSC imaging in mice receiving neoadjuvant (B) or adjuvant (C) therapy. Responders restored CXCR2⁺ MSC recruitment to major organs, indicating alleviation of systemic immunosuppression, whereas non-responders showed persistent loss of recruitment. (D) Longitudinal fluorescence imaging of DiR-labeled CXCR2^+^ MSCs and bioluminescence imaging of metastatic burden in neoadjuvant or adjuvant groups. CXCR2⁺ MSC imaging distinguished responders from non-responders before overt metastatic outgrowth, with earlier prediction in the neoadjuvant group. Notably, multiple mice died with metastases that did not show luciferase signals. FLI: fluorescence imaging; BLI: bioluminescence imaging. *p < 0.05, **p < 0.01, and ***p < 0.001. P values in (B–C) were calculated by comparing the responder and non-responder groups.

In both treatment groups, responders, defined as long-term survivors, progressively alleviated systemic immunosuppression and ultimately restored antitumor immune activity to levels comparable to those in healthy mice. However, the timing of this recovery differed markedly between neoadjuvant and adjuvant treatment. In the neoadjuvant group, systemic immune status became significantly different between responders and nonresponders by day 9 after tumor inoculation (**Fig. 5B, 5D**), 3 days before tumor resection on day 12. In contrast, this distinction did not emerge until day 35 in the adjuvant group (**Fig. 5C, 5D**), 23 days after surgery.

Neoadjuvant chemoimmunotherapy slowed the progression toward systemic immunosuppression before surgery, allowing responders to undergo tumor resection before functional immunosuppression became fully established (**Fig. 5B**). Following surgery, these mice progressively recovered systemic immune activity, ultimately restoring major-organ immune profiles enriched in proinflammatory and antitumor immune populations to levels comparable to those of healthy controls. Notably, the immunosuppression gauge distinguished responders from nonresponders as early as day 9, after three doses of anti-PD-1 and one dose of cisplatin but before surgical resection (**Fig. 5B, 5D**). Thus, CXCR2⁺ MSC imaging identified therapeutic benefit before surgery and before long-term outcomes became apparent. These findings suggest that neoadjuvant chemoimmunotherapy improves postsurgical disease control, at least in part, by limiting the development of systemic immunosuppression before removal of the primary tumor.

In contrast, mice receiving adjuvant therapy had already progressed to established systemic immunosuppression before substantial immune recovery occurred, as indicated by the near-complete loss of CXCR2⁺ MSC recruitment to major organs on days 18 and 23 (**Fig. 5C, 5D**). Responders and nonresponders in this group became distinguishable only after completion of the full treatment regimen, including four doses of anti-PD-1 and two doses of cisplatin (**Fig. 5C, 5D**, day 35). Moreover, responders in the adjuvant group required substantially longer time to restore systemic immune activity to levels comparable to healthy controls than responders receiving neoadjuvant therapy (**Fig. 5B**, day 55 vs. **Fig. 5C**, day 90). These results indicate that adjuvant chemoimmunotherapy can reverse established immunosuppression in a subset of mice, but immune recovery is delayed and less efficient than when treatment is initiated before surgery.

The immunosuppression gauge also provided an early indication of metastatic outcome. Among nonresponders in both treatment groups, IVIS-detectable metastases generally emerged within 7–10 days after the gauge indicated establishment of functional immunosuppression. Some mice developed metastatic lesions without detectable bioluminescence, likely because a subset of metastatic 4T1 cells lost luciferase expression during in vivo progression. Conversely, metastatic burden was not invariably associated with treatment failure. For example, mouse #2 in the adjuvant group developed transient IVIS-detectable metastases (**Fig. 5D**), but these lesions subsequently regressed as the immunosuppression gauge indicated recovery of systemic antitumor immunity. Thus, metastatic burden alone did not fully predict outcome; rather, the ability of the host to prevent or reverse sustained systemic immunosuppression was more closely associated with long-term disease control.

These findings are consistent with emerging clinical evidence that immune checkpoint blockade may be particularly effective when initiated before surgical resection. Neoadjuvant or perioperative immunotherapy trials in early-stage triple-negative breast cancer, including KEYNOTE-522, GeparNuevo, and IMpassion031, have demonstrated improvements in pathological response and/or long-term outcomes, whereas adjuvant-only immunotherapy approaches, including ALEXANDRA/IMpassion030 and A-BRAVE, have not shown comparable improvements in their primary disease-free survival endpoints (37–41). Although these trials were not designed to directly compare neoadjuvant with adjuvant immunotherapy, their collective findings support the importance of treatment timing relative to surgery. Our mechanistic data further suggest that one potential advantage of neoadjuvant therapy is its ability to slow or reverse systemic immunosuppression before it becomes fully established, thereby creating a more favorable immune state at the time of tumor resection.

Overall, longitudinal CXCR2⁺ MSC imaging revealed that therapeutic outcome reflects the interplay among tumor aggressiveness, treatment timing, surgery, and the systemic immune state. Less aggressive 67NR tumors preserved systemic antitumor immunity, allowing tumor resection to achieve durable disease control. In contrast, highly aggressive 4T1 tumors drove progressive systemic immunosuppression that persisted after surgery and was associated with subsequent metastatic outgrowth. Perioperative chemoimmunotherapy improved survival by delaying or reversing this immunosuppressive trajectory, with neoadjuvant treatment producing earlier immune recovery and superior long-term efficacy because intervention occurred before functional immunosuppression was fully established. All the therapy responders ultimately resumed their immune activity comparable to that in healthy mice (**Fig. 4**). These findings support the potential use of the noninvasive immunosuppression gauge to guide treatment selection and timing: surgery may be sufficient for low-risk, less aggressive tumors in immune-active hosts, whereas high-risk, aggressive tumors associated with developing or established systemic immunosuppression may require systemic chemoimmunotherapy, with treatment initiated before surgery offering the greatest opportunity to preserve or restore antitumor immunity and reduce metastatic relapse.

### The immunosuppression gauge captures immune progression across multiple syngeneic and xenograft models

The immunosuppression gauge also demonstrated robust performance across multiple cancer models differing in tumor type and host immune background, supporting the use of CXCL1⁺ neutrophil abundance in major organs as a biomarker of cancer-induced systemic immunosuppression in preclinical studies. Notably, immunocompetent BALB/c mice exhibit a Th2-biased immune phenotype, whereas C57BL/6 mice are relatively Th1-biased. Consistent with these differences, the biodistribution of intravenously administered CXCR2⁺ MSCs differed modestly between the two strains (**Figs. S3, S8**). Similar to BALB/c mice, CXCR2⁺ MSCs were cleared from C57BL/6 mice within one week (**Fig. S9**), enabling repeated longitudinal assessment of immune status in the same animals.

To determine whether the immunosuppression gauge generalizes across tumor types, we next evaluated EO771 breast cancer, LL/2 Lewis lung carcinoma, and B16F10 melanoma, three aggressive syngeneic tumor models previously characterized by pronounced tumor-associated immunosuppression (42–50). Longitudinal CXCR2⁺ MSC imaging revealed a progressive decline in the ability of major organs to recruit CXCR2⁺ MSCs as tumors grew, consistent with the development of systemic immunosuppression. Immune status was already significantly altered when primary tumors reached ∼200 mm³ compared with healthy controls. As tumors progressed beyond their model-specific size thresholds, major organs showed little to no CXCR2⁺ MSC recruitment, indicating the establishment of pronounced systemic immunosuppression (**Figs. 6A– C**).

**Fig. 6.**
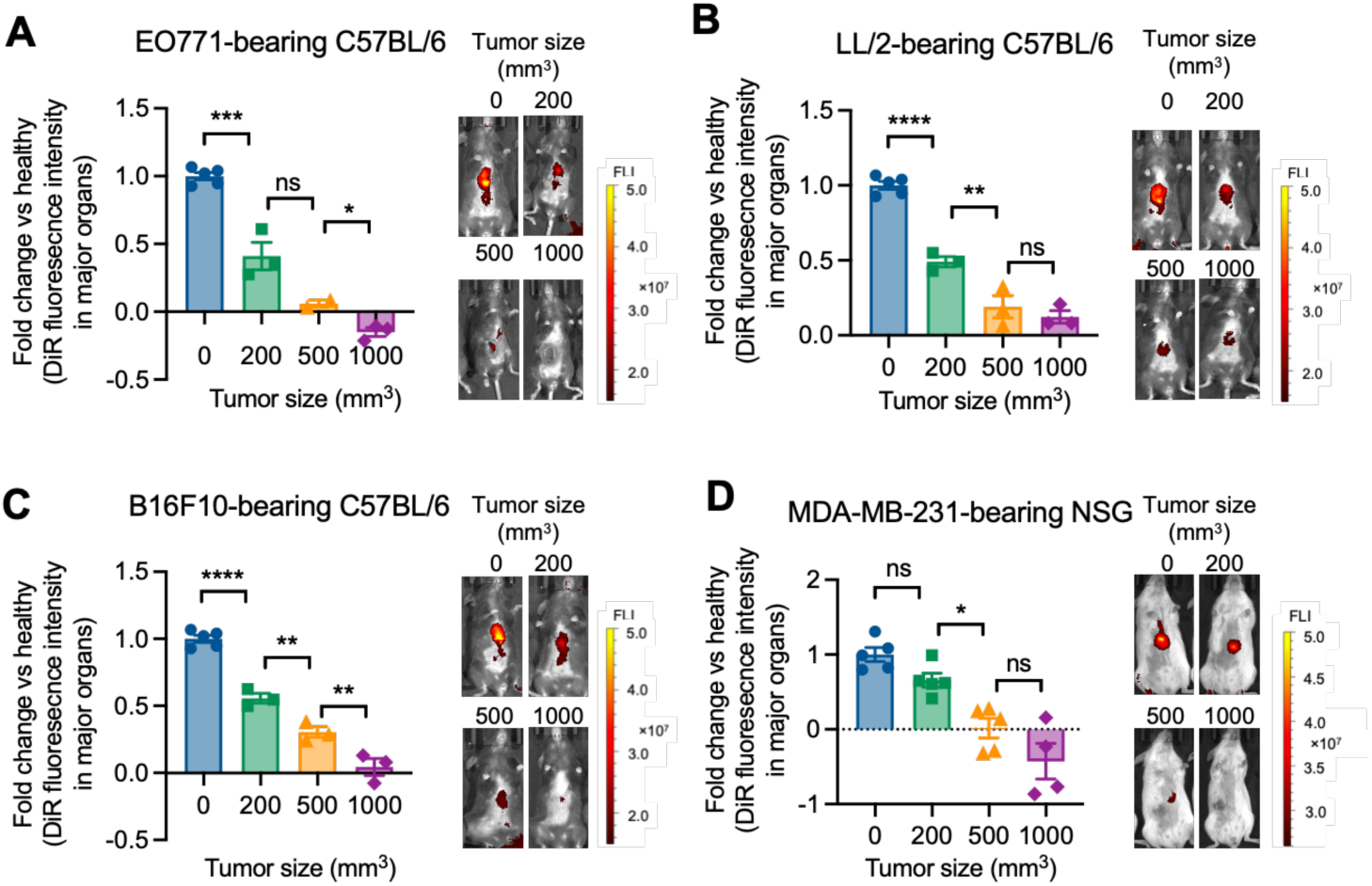
The immunosuppression gauge captures immune progression across syngeneic and xenograft cancer models. (A, B, C) Imaging of DiR-labeled CXCR2⁺ MSCs in syngeneic EO771 breast cancer, B16F10 melanoma, and LL/2 lung carcinoma models established in C57BL/6 mice. (D) Imaging of DiR-labeled CXCR2⁺ MSCs in NSG mice bearing MDA-MB-231 xenograft tumors. The distribution of DiR-labeled cells was assessed by IVIS imaging 24 hours after intravenous injection. Regions of interest (ROI) used to quantify DiR fluorescence in C57BL/6 and NSG mice are shown in **Fig. S8** and **Fig. S10**, respectively. Across all four aggressive cancer models, CXCR2⁺ MSC accumulation in major organs progressively decreased with the growth of aggressive tumors. Mice were imaged by IVIS 24 h after intravenous injection of DiR-labeled CXCR2⁺ MSCs. *p < 0.05, **p < 0.01, ***p < 0.001, and ***p < 0.001.

Interestingly, CXCR2⁺ MSC-based imaging also effectively reflected host immune status in a xenograft model, suggesting that the immunosuppression gauge primarily captures alterations in host myeloid-cell biology rather than adaptive immune responses. MDA-MB-231, an aggressive human breast cancer cell line capable of developing extensive distant metastases at advanced stages (51,52), progressively impaired the recruitment of intravenously administered CXCR2⁺ MSCs to major organs in immunodeficient NSG mice as primary tumors grew (**Fig. 6D**). In contrast to BALB/c and C57BL/6 mice, CXCR2⁺ MSCs predominantly accumulated in the liver of NSG mice (**Fig. S10**). When MDA-MB-231 primary tumors reached ∼500 mm³, major organs showed minimal CXCR2⁺ MSC recruitment, indicating the establishment of pronounced systemic immunosuppression at a stage associated with ongoing metastatic outgrowth (51,52).

Collectively, these findings demonstrate that the CXCR2⁺ cell imaging-based immunosuppression gauge is broadly applicable across diverse preclinical cancer models, including both syngeneic and xenograft systems, and provides a noninvasive approach for longitudinally monitoring dynamic changes in host immune status during cancer progression.

## Discussion

This study harnesses CXCL1⁺ neutrophils as a biomarker and establishes CXCR2⁺ cell trafficking as a noninvasive imaging-based gauge of cancer-induced systemic immunosuppression. Unlike ex vivo T-cell suppression assays, which require terminal tissue collection and primarily detect established functional suppression, this approach enables repeated longitudinal monitoring of immune status within the same host. By tracking the biodistribution of intravenously injected CXCR2⁺ MSCs or healthy donor neutrophils, this gauge quantified late-stage functional immunosuppression comparably to ex vivo T-cell proliferation assays, distinguished tumor aggressiveness at early disease stages, detected elevated metastatic risk before overt metastatic outgrowth, and predicted therapeutic response or resistance during perioperative chemoimmunotherapy. These findings address an important unmet need in cancer research and therapy monitoring: despite the central role of systemic immunosuppression in promoting metastasis and immunotherapy resistance, there is currently no established noninvasive and longitudinally applicable immunosuppression gauge that directly reports the evolving suppressive state of the host.

Application of this immunosuppression gauge across multiple cancer models also advanced our mechanistic understanding of cancer progression and therapy resistance. The results revealed that aggressive tumors induce a staged progression of systemic immunosuppression after tumor inoculation, whereas non-aggressive or poorly migratory tumors, including breast cancer, melanoma, and lung carcinoma models, fail to induce substantial systemic immunosuppression. These distinct tumor–host immune interactions suggest that treatment strategies should be tailored according to both tumor aggressiveness and host immune status. For non-aggressive tumors, surgery alone may be sufficient because preserved systemic immunity limits metastatic outgrowth. In contrast, aggressive tumors may require perioperative chemoimmunotherapy even at early disease stages, before functional suppression is fully established, because surgery alone does not prevent continued progression toward systemic immunosuppression and metastatic relapse. The timing of systemic therapy is also critical. Compared with adjuvant therapy, neoadjuvant chemoimmunotherapy alleviated immunosuppression earlier and more efficiently, allowing hosts to undergo surgery from a less suppressive immune state and thereby improving long-term outcomes. Thus, longitudinal monitoring of host immune status using CXCR2⁺ cell trafficking may help avoid unnecessary systemic therapy in immune-active hosts with non-aggressive tumors, while ensuring timely and sufficient treatment for aggressive tumors that are developing or have established systemic immunosuppression. Notably, all long-term survivors fully alleviated cancer-induced immunosuppression and restored antitumor immunity to levels comparable to those in healthy hosts after therapy. This finding suggests that the gauge may also help determine when treatment has sufficiently restored host immunity and can potentially be discontinued in responders. Beyond distinguishing tumor aggressiveness and predicting therapeutic responsiveness, this imaging-based gauge may therefore provide a practical tool for guiding individualized treatment strategies, assessing immune recovery after therapy, and longitudinally monitoring relapse or metastatic risk in cancer survivors.

A key translational feature of this strategy is that it can be implemented using different CXCR2⁺ cell platforms depending on the experimental or clinical context. CXCR2⁺ MSCs are well suited for preclinical studies because MSCs are easy to expand, genetically engineer, label, and repeatedly administer. MSCs have been widely explored as cellular delivery vehicles for cytokines, growth factors, pro-apoptotic ligands, oncolytic viruses, and other therapeutic payloads in animal models, largely because of their tissue- and tumor-homing properties, low immunogenicity, and ability to release biologically active proteins in vivo (53–55). Receptor engineering can further enhance or redirect MSC homing to desired anatomical sites. For example, prior studies used CXCR2-engineered MSCs to target inflamed mucosa and accelerate tissue repair in inflammatory injury models (56–58). Our study repurposes this chemokine-guided MSC trafficking principle from therapeutic delivery and inflammation targeting to immune-state sensing in cancer. For clinical translation, healthy donor neutrophils may represent a more practical cell source because they naturally express CXCR2 and have existing clinical precedent. Granulocyte transfusions have been used as supportive therapy for severe neutropenia and refractory infections, although their therapeutic efficacy remains debated (59,60). In addition, radiolabeled leukocyte and neutrophil imaging has long been used clinically to detect infection and inflammation, and recent reviews have highlighted the potential of neutrophil imaging in oncology (61,62). Thus, neutrophil-based immunosuppression imaging may face fewer translational barriers than entirely new cell products because it repurposes existing clinical concepts—neutrophil handling, transfusion, labeling, and imaging—for a new application: longitudinal measurement of cancer-associated systemic immunosuppression.

The biological basis of this gauge is the CXCL chemokine–CXCR2 trafficking axis, whose role in metastatic progression is context dependent in major organs that may become metastatic sites. One view is that elevated ELR⁺ CXC chemokines, including CXCL1, CXCL2, CXCL5, and CXCL8, can promote metastatic outgrowth by recruiting CXCR2⁺ myeloid cells, including suppressive MDSCs and pro-tumor neutrophils, and by supporting angiogenesis and pre-metastatic niche formation (63–66). In this context, recruited myeloid cells may mediate functional immunosuppression through mechanisms such as ROS production, PD-L1 expression, IL-10 secretion, and S100A8/A9-associated inflammatory programs (67,68). However, CXCL chemokines are not intrinsically immunosuppressive. They can also recruit inflammatory neutrophils and other immune effector cells that support host defense and antitumor immunity (13). Therefore, the biological consequence of CXCL chemokine activity depends on the phenotype of the recruited CXCR2⁺ cells rather than CXCR2 expression alone.

Our prior study supports this distinction. In breast cancer models, S100A8/A9⁺ neutrophils and CXCL1⁺ neutrophils, defined within the Gr1⁺CD11b⁺Ly6G⁺ myeloid compartment, exhibited immunosuppressive and immune-active phenotypes, respectively (13). CXCL1⁺ neutrophils represented a pro-inflammatory and antitumor myeloid subset, challenging the conventional view that Gr1⁺CD11b⁺Ly6G⁺ myeloid cells uniformly function as suppressive MDSCs. Consistently, immune-active lungs of 67NR-bearing mice showed high CXCR2 expression and abundant CXCL1⁺ neutrophils, but low S100A8/A9⁺ neutrophil abundance (13,69,70). These findings suggest that CXCR2 expression primarily reflects responsiveness to CXCL chemokine gradients and is not, by itself, a strong marker of immunosuppression. In the present study, preserved CXCL1⁺ neutrophil-associated recruitment signals in major organs were associated with reduced systemic immunosuppression and lower metastatic risk, whereas loss of these signals accompanied immunosuppressive progression and metastatic outgrowth. Thus, the prognostic significance of CXCL chemokines in major organs should be interpreted together with the accompanying myeloid-cell phenotype, particularly the balance between CXCL1⁺ neutrophils and S100A8/A9⁺ neutrophils. Notably, clinical analyses further suggest that the CXCL1-to-S100A8/A9 ratio serves as a favorable prognostic biomarker not only in breast cancer but also in lung, colon, prostate, and pan-cancer cohorts (**Fig. S11**) (14).

Several additional issues should be considered in future development. First, the gauge currently reports systemic immunosuppression through the inverse relationship between CXCR2⁺ cell recruitment and organ-level CXCL1⁺ neutrophil-associated signals. Although this relationship was validated across multiple syngeneic and xenograft models, other inflammatory conditions, infections, tissue injuries, or treatment-induced toxicities could also alter CXCR2 ligand gradients and affect cell trafficking (69,70). Future studies should define disease-specific thresholds and incorporate complementary biomarkers to improve specificity. Second, host genetic background and immune bias influence baseline CXCR2⁺ cell distribution, as shown by the differences between BALB/c and C57BL/6 mice. Translation to humans will therefore require establishing individualized or population-level baseline ranges, as well as standardized imaging time points, cell doses, and quantitative organ readouts. Third, although DiR labeling was suitable for preclinical optical imaging, clinical translation would require clinically compatible labeling modalities, such as positron emission tomography (PET), single-photon emission computed tomography (SPECT), or magnetic resonance imaging (MRI)-based cell tracking (71–73). Finally, the mechanistic relationship between CXCL1⁺ neutrophils, antitumor immunity, and metastatic dormancy warrants further investigation. Defining whether these neutrophils directly suppress metastatic outgrowth, recruit effector lymphocytes, reshape myeloid phenotypes, or serve primarily as a surrogate marker of immune-active organs will help determine whether the CXCL1⁺ neutrophil axis is only a biomarker or also a therapeutic target.

In summary, this study introduces a CXCR2⁺ cell-based imaging strategy that functions as a longitudinal gauge of systemic immunosuppression. By converting organ-level CXCL1⁺ neutrophil-associated immune activity into a whole-host imaging readout, this approach detects tumor aggressiveness, metastatic risk, and therapy resistance before overt metastatic outgrowth. The use of engineered CXCR2⁺ MSCs provides a practical platform for preclinical studies, whereas donor neutrophil imaging offers a clinically relevant path for future translation. Together, these findings support immunosuppression-guided cancer management, in which therapy selection and timing are informed not only by tumor size or detectable metastatic burden but also by the evolving systemic immune state of the host.

## Materials and Methods

### Materials

BALB/c mouse mesenchymal stem cells (MSCs) were from Cyagen. EO771 and 4T1/tdTomato/Luc cells were kindly provided by Dr. Lonnie Shea’s lab at University of Michigan. 4T07 and 67NR cells were from Karmanos Cancer Institute at Wayne State University. B16F10, LL/2 and MDA-MB-231 cells were from ATCC. The anti-PD1 antibody (clone RMP1-14; RRID:AB_10949053) was from Bio X Cell. All other antibodies, red blood cell (RBC) lysis buffers, and fixation and permeabilization buffers were from Biolegend. Carboplatin was from Cayman Chemical. Cytokine array membranes were from RayBiotech. Mouse myeloid-derived suppression cell (MACS) isolation kit, mouse naïve CD4^+^ T cell isolation kit, and mouse neutrophil isolation kit were from Miltenyi Biotec. BCA protein assay kit, DiR dye, CellTraceTM Far Red cell proliferation kit, and fetal bovine serum (FBS) were from Thermo Fisher Scientific. Mouse bone marrow mesenchymal stem cell complete medium was purchased from Cyagen. Other cell culture media and supplements were from Gibco. All other chemicals were from Fisher Scientific.

### Cell culture

4T1/tdTomato/Luc, 4T07, and 67NR cells were cultured in high-glucose Dulbecco’s Modified Eagle’s Medium (DMEM) supplemented with 10% FBS, 1% penicillin–streptomycin, and 1% nonessential amino acids. 4T1/tdTomato/Luc cells were additionally maintained with 2 μg/mL puromycin and 300 μg/mL G418. EO771 cells were cultured in high-glucose DMEM supplemented with 10% FBS, 1% penicillin–streptomycin, 10 mM HEPES, 1% nonessential amino acids, and 2 μg/mL puromycin. LL/2 and B16F10 were cultured in high-glucose DMEM supplemented with 10% FBS, 1% penicillin–streptomycin, and 8 μg/mL blasticidin. MDA-MB-231 cells were cultured in high-glucose DMEM supplemented with 10% FBS and 1% penicillin– streptomycin. MSCs were cultured in mouse bone marrow mesenchymal stem cell complete medium. All cells were maintained at 37°C in a humidified incubator with 5% CO₂.

### Stable cell line development

MSCs stably expressing CXCR2 (CXCR2⁺ MSCs) were generated by lentiviral transduction. Briefly, MSCs were seeded in 48-well plates at 3,000 cells/well. After 24 hours, cells were transduced with mCXCR2 lentiviral particles generated from pLV[Exp]-mCherry/Puro-EF1A>mCxcr2[NM_009909.3] at an MOI of 50 in the presence of polybrene. Transduced cells were selected with 2 μg/mL puromycin for 2 weeks to establish a stable CXCR2⁺ MSC line. CXCR2 overexpression was validated by flow cytometry. Briefly, non-engineered MSCs and CXCR2⁺ MSCs were incubated with an FITC-labeled anti-CXCR2 antibody (clone SA044G4; BioLegend; RRID:AB_2566148) for 1 hour, fixed with 4% paraformaldehyde (PFA), and analyzed by a FACSDiscover A8 cell analyzer.

### Healthy donor neutrophil isolation

Lungs from healthy BALB/c mice were collected and processed into single-cell suspensions. Briefly, freshly isolated lung tissues were minced in serum-free culture medium supplemented with 0.1 WU/mL Liberase TL and 150 U/mL DNase I, followed by incubation at 37°C for 30 min. Dissociated tissues were then passed through 70-μm cell strainers to obtain single-cell suspensions. Red blood cells were removed using RBC lysis buffer according to the manufacturer’s instructions. Neutrophils were subsequently isolated from the lung single-cell suspensions by magnetic-activated cell sorting (MACS) using a mouse neutrophil isolation kit according to the manufacturer’s instructions.

### IVIS imaging

CXCR2⁺ MSCs or neutrophils were labeled with the DiR dye according to the manufacturer’s instructions. Briefly, 1 × 10⁶ cells were resuspended in DMEM and incubated with DiR at a final concentration of 10 μM for 20 min at 37°C protected from light. Cells were then washed three to four times with complete culture medium and resuspended in PBS for tail vein injection. CXCR2⁺ MSCs were injected intravenously into BALB/c mice at 5 × 10⁵ cells in 100 μL PBS per mouse. CXCR2⁺ MSCs were injected intravenously into C57BL/6 or NSG mice at 1 × 10⁵ cells in 100 μL PBS per mouse. Neutrophils were injected intravenously into BALB/c mice at 5 × 10⁵ cells in 100 μL PBS per mouse. Whole-body fluorescence imaging was performed using an IVIS spectrum imaging system (PerkinElmer), and images were analyzed using Living Image software. Baseline images were acquired before cell injection to determine background signal and confirm clearance of residual dye signal from prior injections. Mice were then imaged at the indicated time points after injection. Bioluminescence imaging was used to confirm complete resection of 4T1-Luc-tdTomato primary tumors during surgery and to evaluate metastatic burden after surgery. Mice were intraperitoneally injected with 100 μL of 63 mM D-luciferin solution, followed by imaging using the IVIS system.

### Cancer model development

A total of 1 × 10⁶ 4T1-Luc-tdTomato, 4T07, or 67NR cells were injected into the right mammary fat pad of female BALB/c mice. EO771 cells were injected into the right mammary fat pad of female C57BL/6 mice at 1 × 10⁶ cells per mouse. LL/2 or B16F10 cells were injected subcutaneously into the right thigh at 1 × 10⁶ cells per mouse. MDA-MB-231 cells were injected into the right mammary fat pad of female NSG mice at 5 × 10⁶ cells per mouse. Primary tumor size was measured every 2–3 days using calipers and calculated using the formula V = (length × width²)/2.

### Neoadjuvant and adjuvant chemoimmunotherapy strategy

BALB/c mice bearing 4T1/tdTomato/Luc tumors underwent surgical resection on day 12 after tumor inoculation, when tumors in untreated mice reached ∼450 mm³. Neoadjuvant chemoimmunotherapy consisted of 100 mg/kg carboplatin administered on days 6 and 10, together with 100 μg/dose anti-PD-1 antibody (clone RMP1-14; Bio X Cell; RRID:AB_10949053) administered on days 4, 6, 8, and 10 after tumor inoculation. Adjuvant chemoimmunotherapy consisted of 100 mg/kg carboplatin administered on days 18 and 22, together with 100 μg/dose anti-PD-1 antibody administered on days 16, 18, 20, and 22 after tumor inoculation. Both carboplatin and anti-PD-1 antibody were administered intraperitoneally.

### Immune profiling with flow cytometry and cytokine array

Fresh tissues were collected from mice at the indicated time points and processed into single-cell suspensions as described above. For flow cytometric analysis, cells were first blocked with anti-mouse CD16/32 antibodies for 15 min on ice and then stained with an antibody (clone S17011E; BioLegend; RRID:AB_2783138) cocktail targeting cell-surface markers for 30 min on ice. Cells were subsequently fixed and permeabilized, followed by staining with antibody cocktails targeting intracellular proteins at 4°C overnight. After staining, cells were washed with MACS buffer and analyzed using a FACSDiscover A8 cell analyzer. Gating strategies are shown in **Fig. S1.** A total of five antibody panels were used to characterize immune cell populations. Panel 1 was designed to identify CXCL1⁺ neutrophils (CD45⁺CD11b⁺Ly6G⁺CXCL1⁺). This panel included APC-Cy7-conjugated anti-CD45 (clone QA17A26; BioLegend; RRID:AB_2890720), BV711-conjugated anti-CD11b (clone M1/70; BioLegend; RRID:AB_11218791), Pacific Blue-conjugated anti-Gr-1 (clone RB6-8C5; BioLegend; RRID:AB_893559), Alexa Fluor 488-conjugated anti-Ly6G (clone 1A8; BioLegend; RRID:AB_2561340), and Alexa Fluor 647-conjugated anti-CXCL1 (clone 1174A; R&D Systems) antibodies. Panel 2 was designed to identify T helper cells (CD45⁺CD3⁺CD4⁺) and their major subsets, including Th1 cells (IFN-γ⁺) and Th17 cells (IL-17A⁺). This panel included APC-Cy7-conjugated anti-CD45 (clone QA17A26; BioLegend; RRID:AB_2890720), Pacific Blue-conjugated anti-CD3 (clone 17A2; BioLegend; RRID:AB_493644), BV605-conjugated anti-CD4 (clone RM4-5; BioLegend; RRID:AB_11125962), PE-conjugated anti-IFN-γ (clone XMG1.2; BioLegend; RRID:AB_315401), and PE-Cy7-conjugated anti-IL-17A (clone TC11-18H10.1; BioLegend; RRID:AB_2125011) antibodies. Panel 3 was designed to identify regulatory T cells (Tregs), defined as CD45⁺CD3⁺CD4⁺CD25⁺Foxp3⁺ cells. This panel included APC-Cy7-conjugated anti-CD45 (clone QA17A26; BioLegend; RRID:AB_2890720), Pacific Blue-conjugated anti-CD3 (clone 17A2; BioLegend; RRID:AB_493644), BV605-conjugated anti-CD4 (clone RM4-5; BioLegend; RRID:AB_11125962), PE-conjugated anti-CD25 (clone QA19A49; BioLegend; RRID:AB_2894644), and Alexa Fluor 488-conjugated anti-Foxp3 (clone MF-14; BioLegend; RRID:AB_1089114) antibodies. Panel 4 was designed to identify activated cytotoxic T cells (CD45⁺CD3⁺CD8a⁺CD4⁻perforin⁺). This panel included APC-Cy7-conjugated anti-CD45 (clone QA17A26; BioLegend; RRID:AB_2890720), Pacific Blue-conjugated anti-CD3 (clone 17A2; BioLegend; RRID:AB_493644), BV605-conjugated anti-CD4 (clone RM4-5; BioLegend; RRID:AB_11125962), PE-conjugated anti-CD8a (clone QA17A07; BioLegend; RRID:AB_2783127), and FITC-conjugated anti-perforin (clone S16009A; BioLegend; RRID:AB_2910315) antibodies. Panel 5 was designed to identify activated natural killer cells (CD45⁺CD3⁻CD49b⁺perforin⁺). This panel included APC-Cy7-conjugated anti-CD45 (clone QA17A26; BioLegend; RRID:AB_2890720), Pacific Blue-conjugated anti-CD3 (clone 17A2; BioLegend; RRID:AB_493644), PE-conjugated anti-CD49b (clone HMα2; BioLegend; RRID:AB_313029), and FITC-conjugated anti-perforin (clone S16009A; BioLegend; RRID:AB_2910315) antibodies.

For cytokine array analysis, unsorted single-cell suspensions or MACS-isolated cell subsets were cultured in phenol red-free RPMI-1640 medium at a density of 5 × 10⁶ cells/mL at 37°C. Conditioned media were collected after 24 h. Their total protein concentration was quantified using a BCA protein assay kit and then diluted to 500ug/ml for cytokine array study. Cytokine profiling was performed using the Mouse Cytokine Array C3 (RayBiotech) according to the manufacturer’s instructions. Chemiluminescent signals were visualized using an iBright FL1500 Imaging System (Invitrogen) and quantified with iBright Analysis Software (BioRad).

### T-cell suppression ex vivo assay

Lungs and spleens were collected from tumor-bearing BALB/c mice at the indicated time points and processed into single-cell suspensions. Lung-derived Gr1⁺ cell populations were isolated by MACS using a mouse MDSC isolation kit. Naïve CD4⁺ T cells were isolated from spleens by MACS using a naïve CD4⁺ T cell isolation kit and labeled with CellTrace™ Far Red Cell Proliferation Kit according to the manufacturer’s instructions. Gr1⁺ cells and naïve CD4⁺ T cells were resuspended in T cell culture medium consisting of glutamine-containing RPMI-1640 supplemented with 10% FBS, 1% penicillin–streptomycin, 1% nonessential amino acids, 1% sodium pyruvate, and 50 μM 2-mercaptoethanol. Cells were seeded into 96-well plates that had been pre-coated with 2.5 μg/mL anti-mouse CD3ε antibody (clone 145-2C11; BioLegend; RRID:AB_312667) at 4°C overnight. Cells were cultured in the plates at 37°C for 2 h. Three experimental groups were prepared: (1) unstimulated T cells, (2) stimulated T cells, and (3) stimulated T cells co-cultured with lung-derived Gr1⁺ cells at a 1:5 T cell-to-Gr1⁺ cell ratio. T cells were stimulated with 60 μg/mL anti-mouse CD28 antibody (clone 37.51; BioLegend; RRID:AB_312867), 50 U/mL IL-2, and 5 μg/mL anti-mouse CD3ε antibody (clone 145-2C11; BioLegend; RRID:AB_312667). Each condition was prepared in triplicate. Cells were cultured at 37°C for 3 days. After incubation, cells were collected and stained with Brilliant Violet 605™ anti-mouse CD4 antibody (clone RM4-5; BioLegend; RRID:AB_11125962), PE-conjugated anti-mouse CD25 antibody (clone QA19A49; BioLegend; RRID:AB_2894644), and Zombie Green viability dye. Samples were analyzed using a FACSDiscover A8 cell analyzer. CD4⁺ T cells were gated, and T-cell proliferation was assessed based on dilution of the CellTrace Far Red signal.

### Statistics

Data are presented as mean ± standard error of the mean (SEM). The experimental unit and sample size (n) for each analysis are specified in the corresponding figure legends, with n representing biologically independent animals, samples, or experiments, as appropriate. Unless otherwise specified, comparisons between two groups were performed using a two-tailed Student’s t-test while comparison between three or more groups were analyzed using one-way ANOVA. DiR fluorescence signals were compared between two groups using the Mann–Whitney U test. A value of p < 0.05 was considered statistically significant. Statistical analyses were performed using GraphPad Prism version 11.

## Ethics statement

All animal procedures were approved by the Institutional Animal Care and Use Committee (IACUC) at Iowa State University and were conducted in accordance with institutional and applicable federal guidelines.

## Acknowledgments

This work was supported by Roy J. Carver Trust (25-5938) and NIH/NCI R37CA299228-01A1 (JW).

## Authors’ Contributions

YZ conducted all the experiments. JW drafted the manuscript.

