## Supplementary Figures for "An immunosuppression imaging gauge prospectively predicts metastatic progression and immunotherapy resistance in breast cancer"

Step 1: gate CD45<sup>+</sup> immune cell population

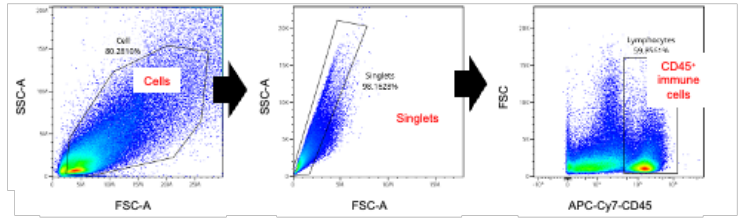

Step 2: gate immune cell subsets

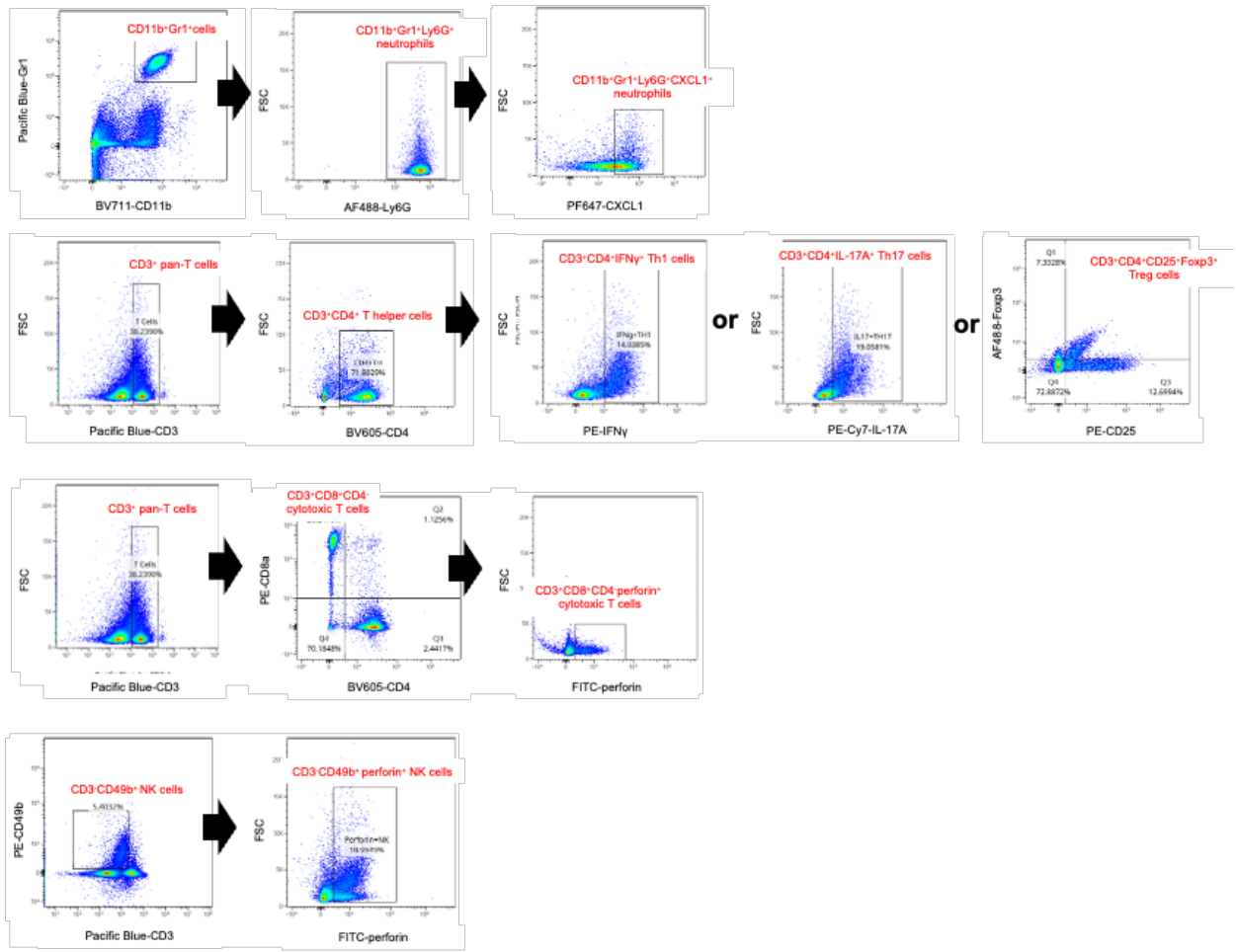

**Fig S1. Gating strategy for flow cytometric analysis of immune cell populations.** Representative gating strategy for CXCL1<sup>+</sup> neutrophils, T helper cell subsets, and activated effector cells, including perforin<sup>+</sup> cytotoxic T cells and perforin<sup>+</sup> NK cells. NK: natural killer.

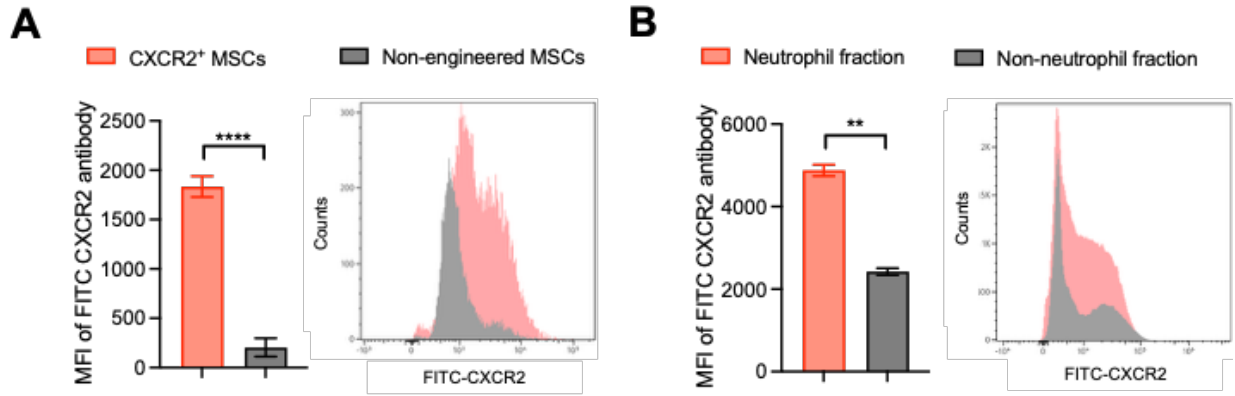

**Fig S2. Surface expression of CXCR2 on CXCR2<sup>+</sup> MSCs and healthy donor neutrophils.** CXCR2 expression on the surface of (A) CXCR2<sup>+</sup> MSCs and non-engineered MSCs (control), and (B) neutrophil and non-neutrophil fractions isolated from healthy mouse lungs by magnetic-activated cell sorting. Cells were stained with FITC-conjugated anti-mouse CXCR2 antibody and analyzed by flow cytometry. MFI: mean fluorescence intensity. \*\*p < 0.01, and \*\*\*\*p < 0.0001.

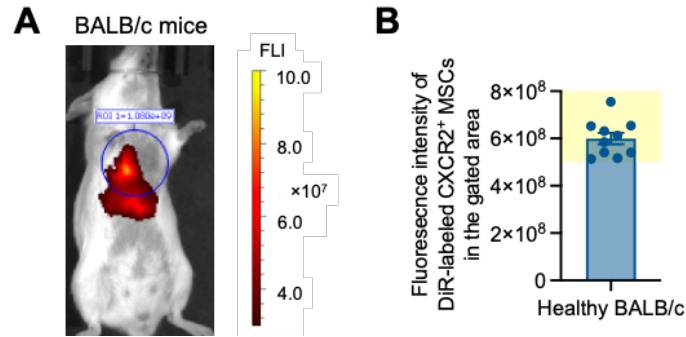

**Fig S3. ROI and baseline distribution of DiR-labeled CXCR2<sup>+</sup> MSCs in BALB/c mice.** (A) Region of interest (ROI) used to quantify DiR-labeled CXCR2<sup>+</sup> MSC accumulation in major organs and (B) the corresponding range of DiR signals measured in healthy BALB/c mice. The yellow-shaded area in (B) indicates the range of DiR fluorescence intensities measured in healthy BALB/c mice.

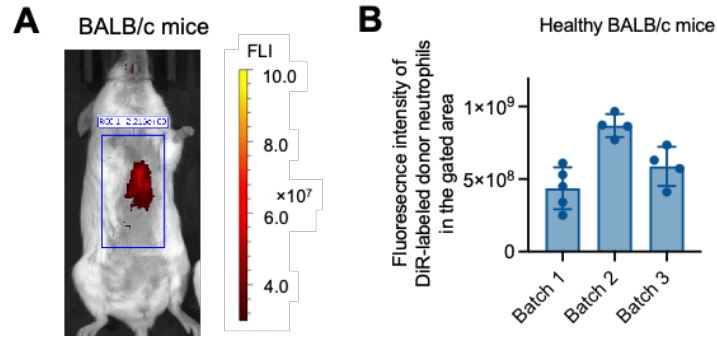

**Fig. S4. ROI and baseline distribution of DiR-labeled healthy donor neutrophils in BALB/c mice.** (A) ROI used to quantify DiR-labeled healthy donor neutrophil accumulation in major organs and (B) the corresponding range of DiR signals measured in healthy BALB/c mice using neutrophils collected from different donor mice. Three batches of healthy donor neutrophils were used to image healthy BALB/c mice. Because neutrophils collected from different healthy donors or at different time points exhibited variable accumulation in major organs, healthy mice were included as controls in each experiment and received neutrophils from the same batch administered to tumor-bearing mice. Organ accumulation in tumor-bearing mice was then normalized to that in the corresponding healthy controls to quantify the degree of systemic immunosuppression.

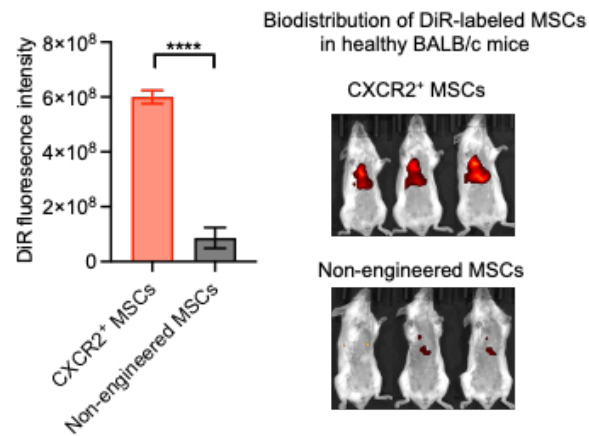

**Fig S5. Distribution of DiR-labeled non-engineered MSCs and CXCR2<sup>+</sup> MSCs in healthy BALB/c mice.** Non-engineered MSCs cannot home to major organs abundant with CXCR2 ligands. \*\*\*\*p < 0.0001.

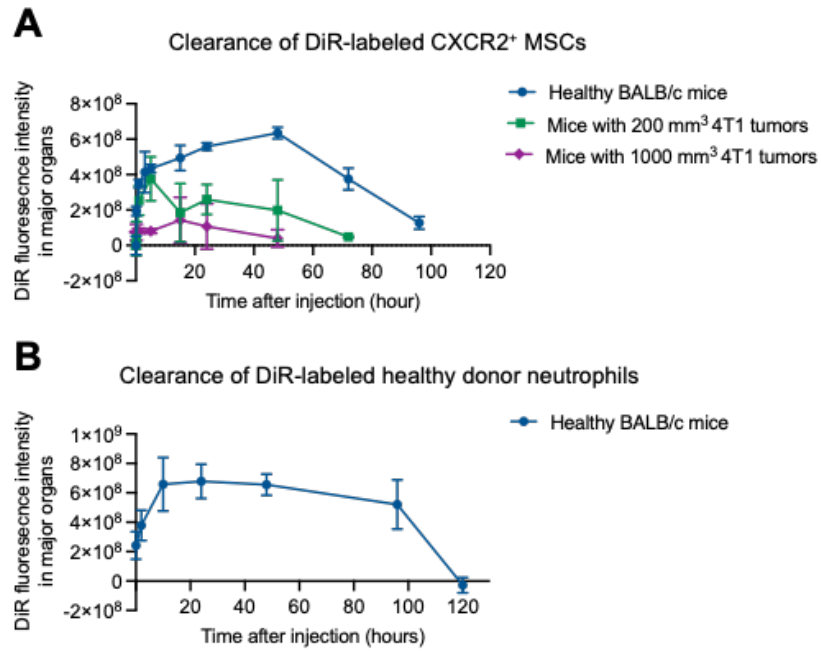

**Fig S6. Clearance kinetics of DiR-labeled CXCR2<sup>+</sup> MSCs and healthy donor neutrophils from healthy BALB/c mice or mice bearing 67NR, 4T07, or 4T1 tumors.** Mice were imaged by IVIS at desired time points after intravenous injection of DiR-labeled cells.

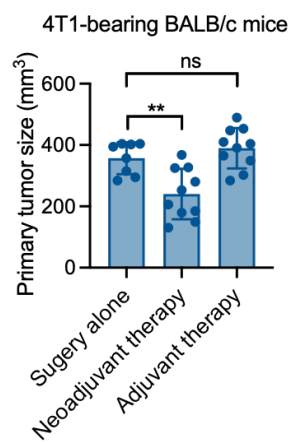

**Fig. S7. Primary tumor size at the time of surgical resection in 4T1-bearing mice receiving surgery alone, neoadjuvant chemoimmunotherapy, or adjuvant chemoimmunotherapy.** Tumor resection was performed on day 12 after tumor inoculation.

C57BL/6 mice

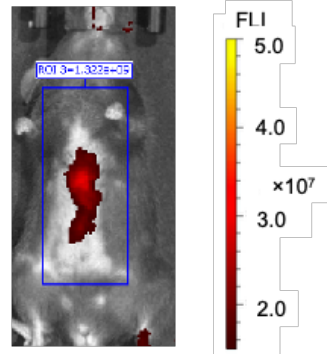

**Fig S8. ROI of DiR-labeled CXCR2<sup>+</sup> MSCs in C57BL/6 mice.** Region of interest (ROI) used to quantify DiR-labeled CXCR2<sup>+</sup> MSC accumulation in major organs in healthy C57BL/6 mice.

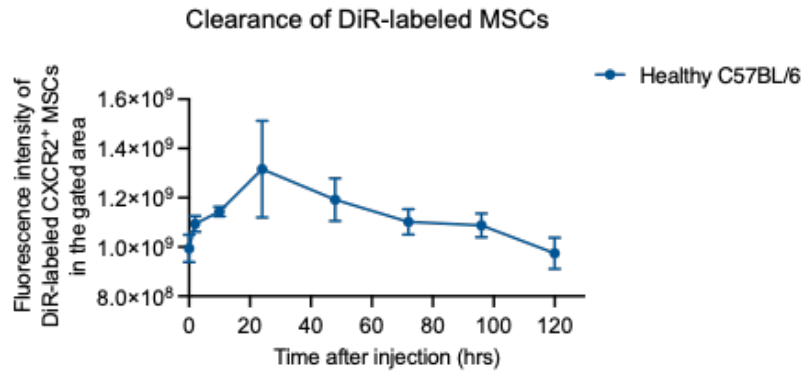

**Fig. S9. Clearance kinetics of DiR-labeled CXCR2<sup>+</sup> MSCs from C57BL/6 mice.** Mice were imaged by IVIS at desired time points after intravenous injection of DiR-labeled CXCR2<sup>+</sup> MSCs.

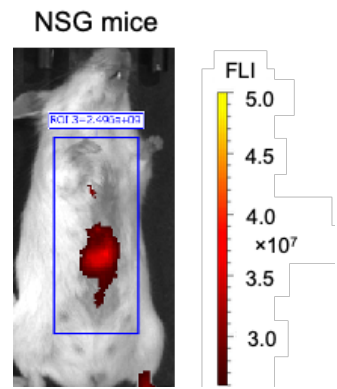

**Fig S10. ROI of DiR-labeled CXCR2<sup>+</sup> MSCs in NSG mice.** Region of interest (ROI) used to quantify DiR-labeled CXCR2<sup>+</sup> MSC accumulation major organs in healthy NSG mice.

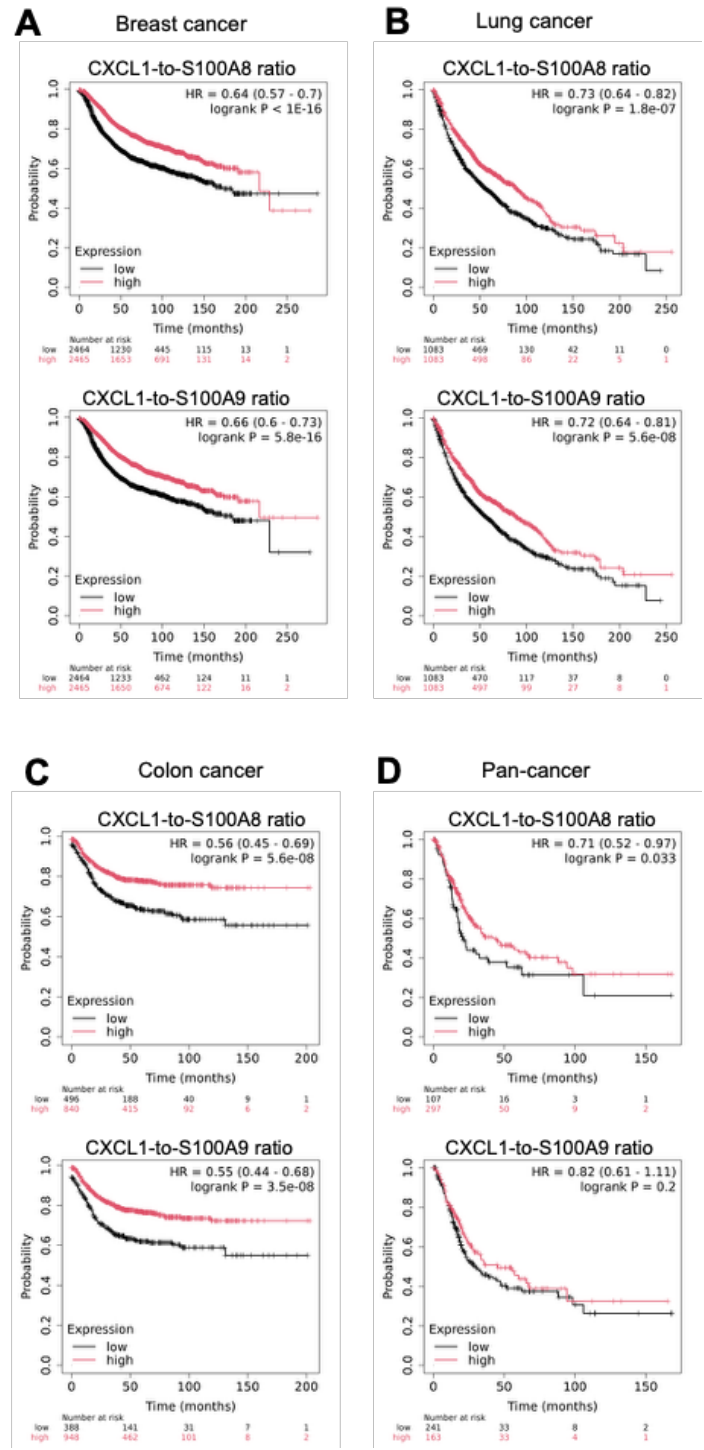

**Fig S11. The CXCL1-to-S100A8/A9 ratio is a favorable prognostic biomarker across multiple clinical cancer cohorts.** Kaplan-Meier survival plots of breast (A), lung (B), colon (C), and pan-cancer patients (D) stratified by high or low CXCL1-to-S100A8/A9 ratios. Data were obtained from KMplot.com.
